# Human-in-the-Loop Weight Compensation in Upper Limb Wearable Robots Towards Total Muscles’ Effort Minimization

**DOI:** 10.1101/2020.11.02.366070

**Authors:** Rezvan Nasiri, Hamidreza Aftabi, Majid Nili Ahmadabadi

## Abstract

In this paper: (1) We present a novel human-in-the-loop adaptation method for whole arm muscles’ effort minimization by means of weight compensation in the face of an object with an unknown mass. (2) This adaptation rule can also be used as a cognitive model for the identification of mass value using EMG sensors. (3) This adaptation rule utilizes the activation (myoelectric) signal of only four muscles in the upper limb to minimize the whole muscles’ effort. We analytically discuss the stability, optimality, and convergence of the proposed method. The effectiveness of this method for whole muscles’ effort reduction is studied by simulations (OpenSim software) on a generic and realistic model of the human arm, a model with 7-DOF and 50 Hill-type-muscles. The simulation results show the presented method’s performance and applicability for weight compensation and mass estimation in upper limb assistive robots. In addition, the simulations in OpenSim completely support that the suggested set of mono-articular muscles are sufficient for whole muscles’ effort reduction.

## I. Introduction

Despite of many efforts in design and control of upper limb wearable robots [1], [2], still, design of an effective control approach for upper limb assistance is a challenge; see [3-6]. A proper control approach should consider the human as a part of the control strategy. Consequently, several considerations should be met; the controller should: (1) minimize muscles’ effort/force, (2) not impose a trajectory to the human, (3) utilize biofeedback, and (4) be adaptive.

An effective approach for upper limb assistance without imposing a reference trajectory is weight compensation, which is a task-independent (trajectory free) method; i.e., it is applicable for both cyclic and noncyclic tasks. Since the upper limb exoskeletons are mostly designed for picking, carrying, and placing heavy objects with slow dynamics, the gravitational torque is a high portion of required torque at each joint; i.e., weight compensation can reduce the muscle effort/force drastically. There are two different methods for weight compensation active and passive. The former generates the gravitational torque by actuators [7-10] and the later compensates the gravity effects with passive elements; e.g., spring or counterbalance [11-13]. However, most of the weight compensation exoskeletons are designed to generate a predefined torque profile; they are not useful for unknown object weight compensation. Besides, human biomechanics has a time-varying nature even in repetitive tasks (e.g., walking) [14-16], which makes having an adaptive strategy to be a must [17].

There are many adaptive control approaches that update the exoskeleton torque based on the instantaneous feedback of the human body; see [6], [18], [19]. Among all possible biofeedback options, the myoelectric signal is one of the best choices for torque adaptation due to its fast dynamics and monotonic relation with muscle force [20], [21]. At first glance, the myoelectric signal seems to be a noisy signal; however, by using an appropriate signal processing it provides us with more biomechanical information compared to force and motion sensors; it can be used to estimate muscle force [22], fatigue [23], metabolic rate [24], and even discharged timing of motor-neurons [25]. In addition, thanks to recent technology developments, myoelectric signals (e.g., EMG) are considered as accurate, cost-efficient, and small-sized sensors [26] such that the recent commercialized prosthetic hands are benefiting them for real-time motion control [27].

There are many works benefit the EMG sensors for upper limb exoskeleton torque adaptation; e.g., see [4-6], [28-31]. Nevertheless, the proposed methods require the EMG signal of all contributing muscles which make them impractical, very complex, expensive, and time-costly; e.g., [28], [29] utilizes 16 EMG sensors. There are also some other works that utilized a smaller number of EMG sensors; see [4], [6], [30], [31]. However, these works are restricted to a single joint (mostly elbow), and they are basically designed for a specific task or device. In another perspective, they did not suggest a general and optimal adaptation rule with a minimum number of sensors to achieve “total muscles’ effort reduction.”

Recently in [32], we presented the hypothesis that “feed-back from mono-articular muscles is sufficient for exoskeleton torque adaptation and whole muscles’ effort reduction.” Benefiting from this hypothesis, we present a novel, simple, and effective weight compensation approach using a minimum number of EMG sensors, a total of 4 EMG sensors for the whole arm. This generic adaptation method does not limit the users’ voluntary motions, is task-independent (can be used for both cyclic and noncyclic tasks), and can be applied to different types of upper limb assistive devices for weight compensation. In addition, this adaptation rule provides us with a cognitive model to identify the object’s mass value, which can be used as a model to describe human understanding about the environment by sensory information of muscles’ force and joint positions. In recent years, the development of reliable and realistic simulation toolboxes as OpenSim [33] draws much attention to the model-based analysis of human biomechanics. Hence, in this paper, the developed method is analyzed using a generic model in OpenSim with 7-DOF and 50 Hill-type-muscles.

## II. Problem Statement

Consider the dynamical equations of the human-arm augmented by an upper-limb-exoskeleton (see Fig.1) as^1^:

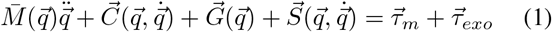

where 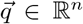 is the vector of the joints’ positions, *n* is number of joints and *m* is the number of all contributing muscles. 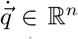 and 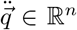 are first and second ordered time derivative of 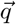. In addition, 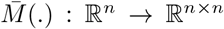 is the mass matrix and 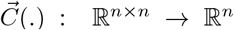 is the vector of Coriolis and Centrifugal force. 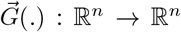is the vector of gravity force. And,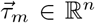 is the muscles’ net torque which is a summation of muscles’ force; i.e., for *j*th joint we have 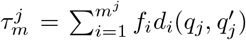 where *f*_*i*_ *∈* ℝ^+^ and *d*_*i*_ *∈* ℝ are *i*th muscles’ force and lever arm, and *m*_*j*_ is the number of contributor muscles’ at *j*th joint. Since muscles are either mono-or bi-articular *d*_*i*_ is a function ofboth targeted joint (*q*_*j*_) and its adjacent joint 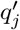 positions. 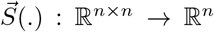 is added to model dynamical effectsof other biological elements; e.g., ligaments and tendons. Finally, 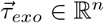 is the torque applied by exoskeleton^2^.

Assuming that the human picks, carries, or places an object with unknown mass (*m*^*\**^ *∈* ℝ^+^) and the exoskeleton is trying to compensate the effects of unknown mass by applying an external torque 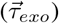, the dynamical equations in Eq.1are rewritten as:

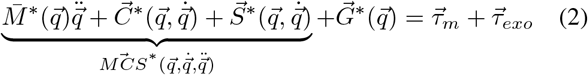

where 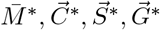 are parameters which are defined in Eq. 1 and affected by *m*^*\**^ *>* 0; if 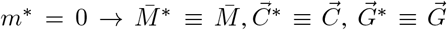, and 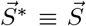. In this equation, 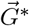 can be divided in two terms as 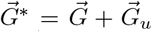 where 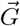 is the gravity vector for *m*^*\**^ = 0 and 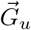 is the gravitational effects of *m*^*\**^. 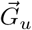 can be written as a function of position 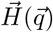 multiplied by *m*^*\**^*g* as 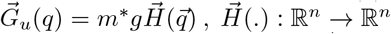 where *g* = 9.81*m/s*^2^ is the gravity acceleration and 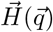 is a position dependent vector. Hence, Eq.2is presented as:

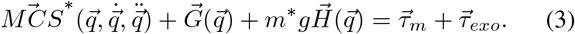

To compensate the gravity effect of an object, simply we can cancel out its dynamical effect by setting 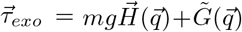, where the first term is an adaptive term and, *m ∈* ℝ is left to adaptation, and 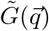 is the estimation of 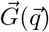. In addition, 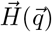 is defined based on the exoskeleton kinematics; it is a predetermined function of position.

Henceforth, the problem is to present an adaptation method for *m* and updating the exoskeleton torque 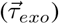 to minimize the total muscles’ effort. The total muscles’ effort is defined as 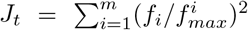 where 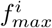 is the *i*th muscle’s maximum isometric force [34]. Fig. 1 shows schematics for this adaptation method in which the adaptation rule block uses the EMG signal as biofeedback.

**Fig. 1.**
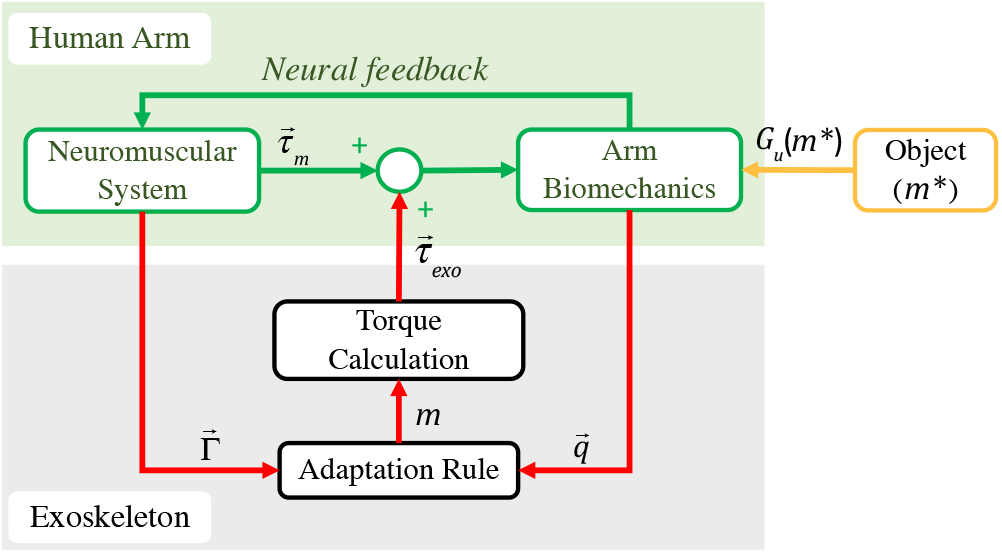
The human-in-the-loop control block diagram: Human arm in interaction with an object with unknown mass (*m\**) and weight compensation exoskeleton system; *Gu* is the gravitational effects of object. The adaptation rule utilizes the EMG signal of chosen muscles along with the joint positions to estimate mono-articular muscles’ torque (*τmm*) and adapt *m*. The torque calculator block uses the adapted value of *m* to update the exoskeleton torque.

The adaptation of *m* for minimization of *J*_*t*_ does not necessarily converge to *m*^*\**^. To **(1)** minimize *J*_*t*_ and **(2)** identify *m*^*\**^ at the same time by *m* adaptation, some certain conditions should be met. In the following section, we discuss this point with more details.

## III. Mathematical Analysis

Whole of our mathematical analysis are presented without any assumption on the shape of the motions (cyclic or noncyclic) and the exoskeleton dynamics.

### 1) Optimality

As we proved in [32], the “whole muscles’ effort” (*J*_*t*_) reduction by exoskeleton torque optimization 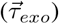 is equivalent with the “squared of two antagonistic mono-articular muscles’ torque” minimization (*J*_*mm*_). In other words, the gradient of *J*_*t*_ w.r.t. the exoskeleton torque is in the same direction of the gradient of *J*_*mm*_ w.r.t. the exoskeleton torque; i.e., 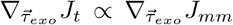. Hence, to extract the adaptation, we use *J*_*mm*_ instead of *J*_*t*_ as:

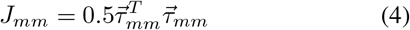

where 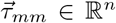 is the torque vector of two antagonistic mono-articular muscles at each joint; for *n* joints, 2*n* mono-articular muscles are required. The muscle net torque can be rewritten as 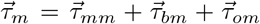 where 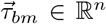 is the torque vector of bi-articular muscles and 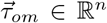 is the torque vector of remained mono-articular muscles; i.e.,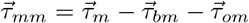.

### 2) Adaptation rule

To extract the adaptation rule, we substitute 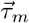 with 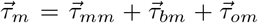 and 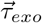 with 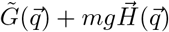 in Eq.3, and compute 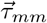 as follows.

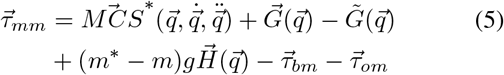

By applying the gradient descent on *J*_*mm*_ as:

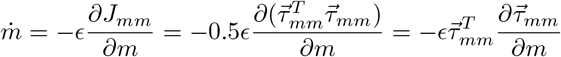

and using Eq.5, the adaptation rule is computed as:

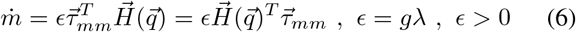

Where ϵ is the adaptation rate which can control the adaptation performance; i.e., speed and accuracy of convergence. Using Eq. 6 leads to instantaneous *Jmm* minimization, and based on hypothesis presented in [32], consequently, minimization of *J*_*mm*_ leads to *J*_*t*_ reduction. Nevertheless, 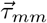 cannot be directly computed, and in practice we estimate 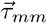 for each joint using myoelectric signal (EMG) of monoarticular muscles as 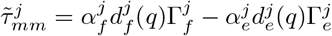 where *α, d*(*q*), and Γ are tunning scale, muscle lever, and RMS EMG at *j*th joint, respectively; see [32] and [35]. Besides, *f* and *e* indicates values related to flexor and extensor muscles. Due to imperfect estimation of 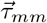, we have 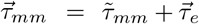 where 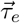 is error torque which is added to model imperfections of our estimation such that 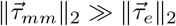.

#### 3) Stability & convergence

To prove the stability and convergence of the adaptation rule (Eq. 6), we utilize 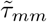 instead of 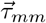 in the adaptation rule (Eq.6), use Eq.5, and rewrite the adaptation dynamics as:

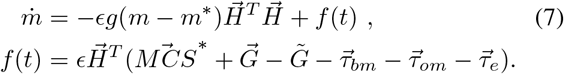

##### Theorem 1

(Stability & Convergence). The adaptation dynamics (*Eq.7):*

*1) is stable*.

*2) is convergent towards m*^*\**^ *with error of δm, if (****C-1****) motions are sufficiently slow* 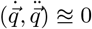.

*3) is convergent towards m*^*\**^, *if (****C-1****) motions are sufficiently slow* 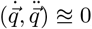, *and (****C-2***) 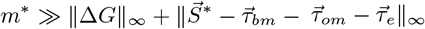.

*Proof. Stability proof:* The dynamical equation in Eq.7is a summation of two terms 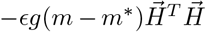 which is a convergent term towards *m* = *m*^*\**^ **(2)** *f* (*t*) which is a non-vanishing perturbation. Based on [36, pp.346], the overall adaptation dynamics (Eq.6) is ultimately bounded and stable if and only if the first term is globally asymptotically stable and the second term is bounded.

To prove 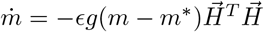 is globally asymptotically stable and convergent towards *m* = *m*^*\**^, we choose *V* = 0.5(*m* − *m*^*\**^)^2^ as a Lyapanov candidate. The time derivative of *V* is 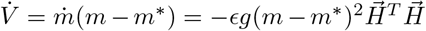 where 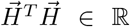 is a semi-positive definite multiplier, which makes 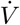 semi-negative definite. Hence, based on LaSalle’s invariance principle presented in [36, pp.128], 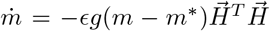 is globally asymptotically stable dynamics convergent to *m* = *m*^*\**^.

*f* (*t*) is a summation of sufficiently smooth functions of arm position, velocity, and acceleration. In addition, in human motions, 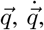, and 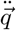 are also sufficiently smooth and bounded functions of time; thus, *f* (*t*) is a bounded disturbance for Eq.7. Therefore, the overall system is stable. Note that the stability of Eq.7is proved without any assumption on the dynamics.

*Convergence proof:* If we assume that 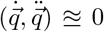, consequently it is concluded that 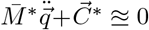. In this case, the adaptation dynamics is 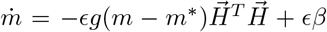 where 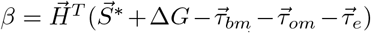 is almost a fixed function of time and 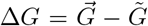 is the gravity compensation error. Hence, the adaptation dynamics is rewritten as 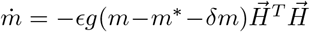 where 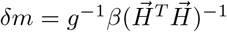 is mass convergence error.

If the dynamical effects of unknown mass is dominant to gravity compensation error (Δ*G*) and the dynamical effects of the biological elements 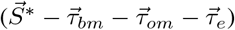 as *m*^*\**^ *≪* | |*δm*| | _*∞*_ or 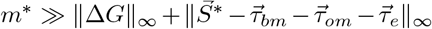 (**C-2**) the adaptation dynamics is 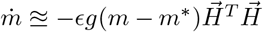which is convergent to *m*^*\**^. If the conditions are not satisfied *m* converges to *m*^#^ = *m*^*\**^ + *δm*.

The adaptation rule is extracted to minimize *J*_*mm*_; it is a gradient descent over *J*_*mm*_. Hence, the equilibrium point of adaptation dynamics (*m*^#^) minimizes the *J*_*mm*_ and consequently based on [32] minimizes the total muscles’ effort (*J*_*t*_). However, *m*^#^ does not necessarily identify *m*^*\**^; in general *m*^#^ = *m*^*\**^ + *δm*. Nevertheless, if **C-1** and **C-2** are satisfied, we can identify *m*^*\**^ along with *J*_*t*_ minimization; i.e., *δm ≊* 0 → *m*^*\**^ *≊m*^#^. **C-2** indicates that Δ*G ≊* 0; i.e., the exoskeleton weight should be known properly which is not a challenge. In the following simulations, we study the performance of adaptation rule (Eq.6) on *J*_*t*_ reduction and the deviations of *m*^#^ from *m*^*\**^ in cases that **C-1** is violated.

## IV. Simulation

In this section, using the OpenSim software [37] with MATLAB API [38], we study our adaptation rule’s performance in three different tasks; in these tasks, we violate **C-1**. The musculoskeletal model utilized in this simulation is presented by Holzbaur *et al*. [39], which is a modified version of the previously released one [40]. This upper extremity model contains 7 degrees of freedom (shoulder 3-DOF, elbow 1-DOF, and wrist 3-DOF) and 50 Hill-type muscles introduced by Zajac *et al*. [41]. We scaled the generic model using the dataset presented in [39] to reach a model consistent with human anthropometry.

The block diagram of our approach for running MATLAB-OpenSim simulation is presented in Fig.2 where the adaption rule is realized in MATLAB and arm biomechanics is modeled by OpenSim. The simulation is chopped for 168 time steps in 37*s*; each time interval is equal to 220*ms*. At each interval, the joint position and muscle forces are computed using OpenSim and feed to MATLAB, where MATLAB updates the exoskeleton torques using the adaptation rule. OpenSim calculates the muscles’ forces using the Computed Muscle Control (CMC) algorithm [42]. The CMC algorithm includes three parts; PD controller, static optimization block, and forward dynamics block. In the first step, the CMC algorithm utilizes a PD controller to track reference kinematics (the designed task for the arm) by computing the desired accelerations. In the next step (static optimization in Fig. 2), CMC finds muscle forces by solving the following optimization problem [42]:

**Fig. 2.**
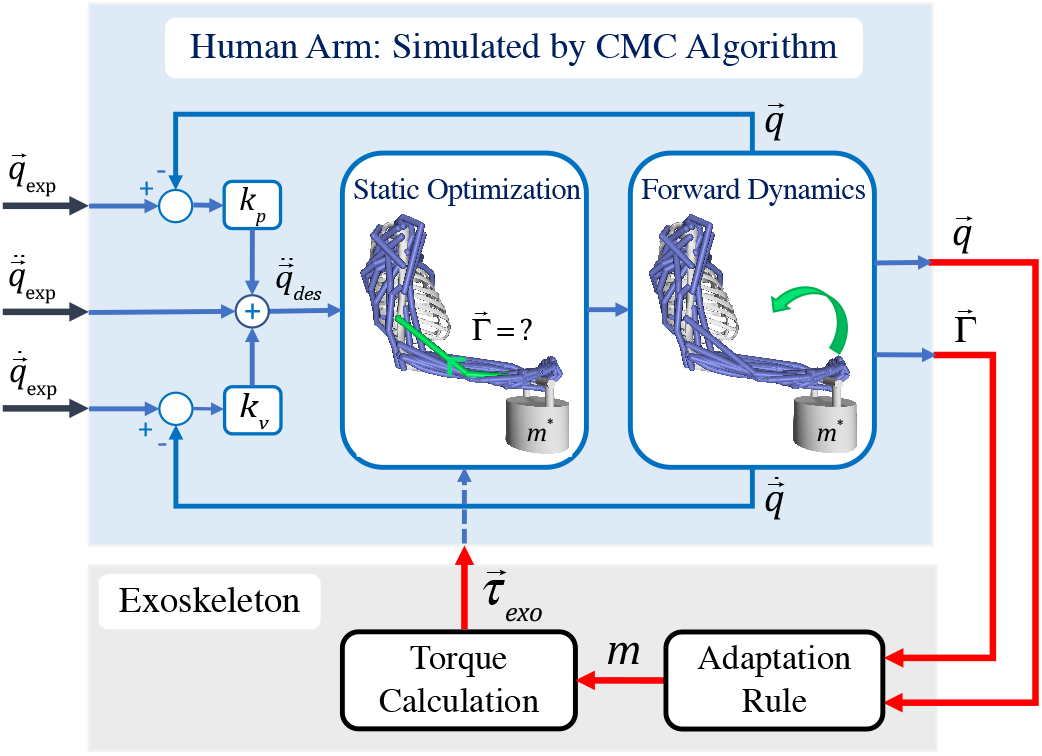
Closed-loop control of human upper limb model augmented with exoskeleton in OpenSim software. In the first step, the CMC algorithm, which includes a PD controller, static optimization block, and forward dynamics block, finds muscle forces. In the next step, the mass estimation block uses Eq.6, joints’ position, and the force signal of four suggested mono-articular muscles to estimate the mass of an unknown object. Finally, the torque calculation block uses *m* to calculate the assistive torque.

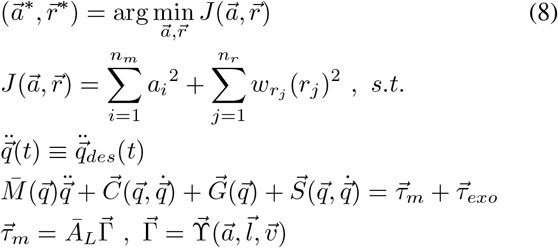

where *a* is muscle activation, *n*_*m*_ is the number of muscles, *r* is the reserve actuator’s torque, *n*_*r*_ is the number of reserve actuators, and 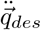 is the desired acceleration (the output of PD controller in Fig. 2).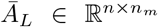 is the lever arm matrix, and 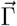is the vector of muscle forces which is computed using Hill-model function 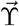; in Hill-type muscles, the muscle force (*f*_*i*_) is a function of muscle’s activation (*a*_*i*_), muscle’s length (*l*_*i*_), and muscle’s velocity (*v*_*i*_) [41]. It is important to note that reserve actuators are added to each coordinate to make up for strength deficiencies in muscles and enable the simulation to run. However, their weights (*w*_*r*_) are chosen big enough; thus, using reserve actuators is highly penalized in the optimization, and the generated forces by reserve actuators are all within the best possible boundaries.

To consider the unknown mass, a bucket with a mass of 5*kg* is attached to the model hand; see Fig.3a. The augmented exoskeleton in the upper limb model is designed as two 1-DOF ideal actuators at the elbow and shoulder joints, which apply assistive torque at the sagittal plane 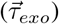. The task in our simulation is to move the bucket on a circular trajectory in the sagittal plane with the radius of 12*cm* and three different frequencies; *ω* = 1*rad/s*, 2*rad/s*, and 4*rad/s*. The main purpose of this task definition is to study the whole muscles’ effort reduction and the equilibrium point of adaptation dynamics for **C-1** violation; 4*rad/s* is an extreme violation.

*m* is estimated using Eq.6, joints’ position and the force signal of four suggested mono-articular muscles near to the skin; Biceps and Triceps as antagonistic mono-articular muscles in the elbow joint and Pectoralis major and Latissimus dorsi as antagonistic mono-articular muscles in the shoulder joint. The estimated *m* is used to calculate the assistive torque 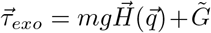 where 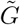 is the estimated gravity vector of the musculoskeletal model.

To study the convergence behavior of the adaptation for each frequency, the simulation is run from two different initial masses; *m*_0_ = 1*Kg* and *m*_0_ = 9*Kg*. The results of this simulation are presented in Fig.3. Fig.3b shows the estimated mass value in the course of adaptation. Interestingly, the equilibrium point (*m*^#^) for each frequency is identical regardless of the initial point. Clearly, by moving from *ω* = 1*rad/s* to *ω* = 4*rad/s*, the deviations of equilibrium point (*m*^#^) from *m*^*\**^ = 5*Kg* increases with a linear pattern such that *∀ ω ∈* [0 4] → *m*^*\**^ *≊m*^#^ + 0.38*ω* with estimation confidence of 95%.

**Fig 3.**
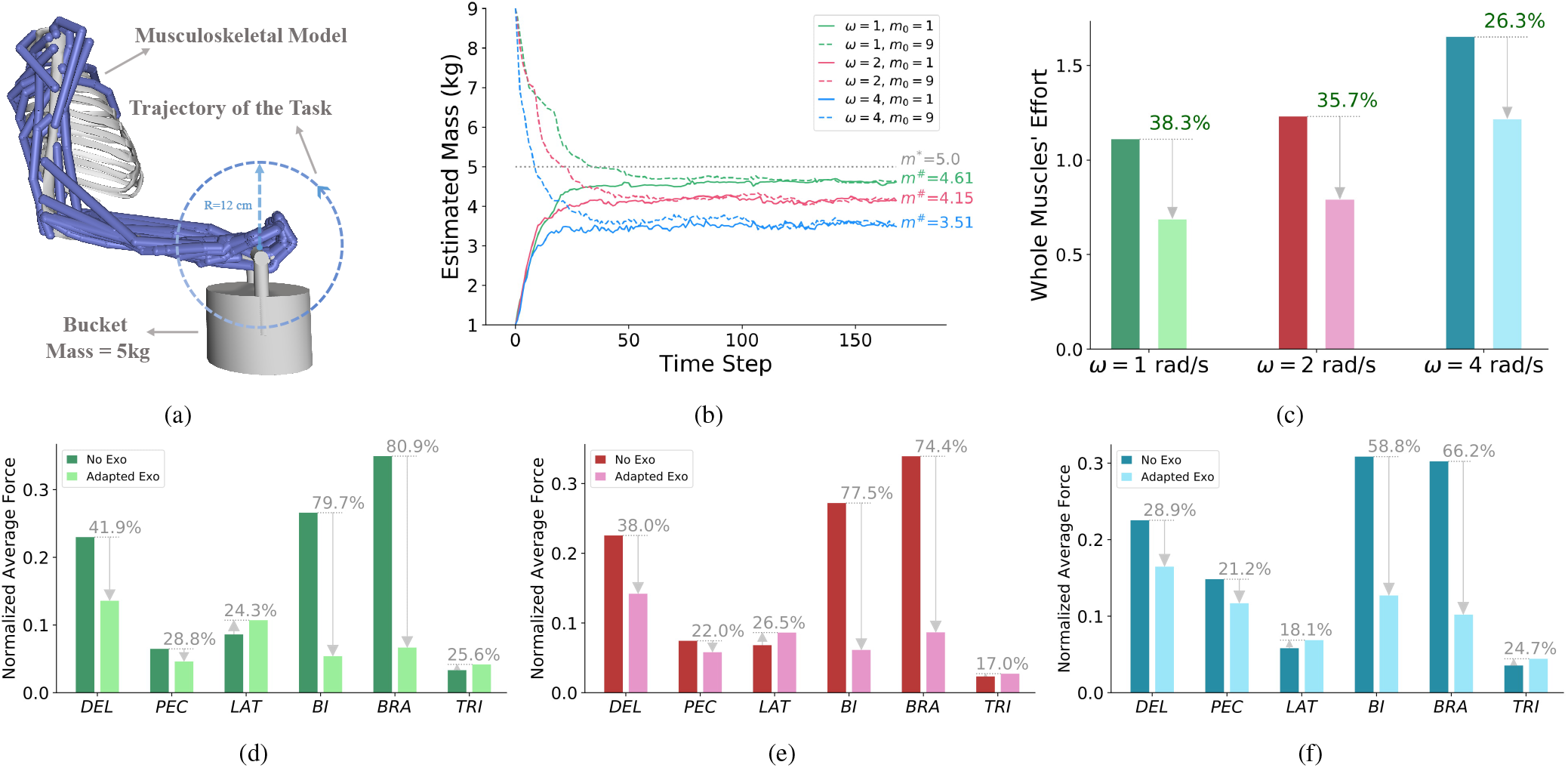
(a) is the utilized biomechanical model in OpenSim for analyzing the behavior of adaptation in different conditions. The model consists of 7-DOF (shoulder 3-DOF, elbow 1-DOF, and wrist 3-DOF) and 50 Hill-type muscles. The task is to move the object with an unknown mass of *m*^*\**^ = 5*Kg* over a circular trajectory with a radius of 12*cm* and three different frequencies *ω* = 1*rad/s*, 2*rad/s*, 4*rad/s*. The exoskeleton consists of two 1-DOF ideal actuators that apply torque at shoulder and elbow joints in the sagittal plane. The goal is to compensate for the gravity effect of unknown mass at elbow and shoulder joints using force feedback (EMG signal) from four mono-articular muscles at elbow and shoulder joints. (b-f) are OpenSim simulation results for simultaneous mass identification and assistive torque adaptation. (b) shows the adaptation performance for mass identification when the initial mass is set on *m*_0_ = 1*Kg* and *m*_0_ = 9*Kg*, respectively. As it is clear, regardless of the initial point, the converged mass value is identical for each frequency. And, by increasing the frequency of motion, the converged mass value decreases from *m*^*\**^. (c) shows the whole muscles’ effort cost function (*Jt*) before and after adaptation. Clearly, the adaptation is successful for the whole muscles’ effort reduction. (d-f) compare the forces of six major muscles before and after adaptation in three different frequencies, respectively; Deltoid (DEL), Pectoralis major (PEC), Latissimus dorsi (LAT), Biceps (BI), Brachialis (BRA), and Triceps (TRI). Based on these figures, the forces of the two muscles are increased. Note that reducing the main cost function (the average efforts of all muscles) does not necessarily mean each individual’s effort reduction.

Fig.3c-f describes the effect of *m* adaptation on total muscles’ effort (*J*_*t*_) and the force of six major muscles in the arm. Based on Fig.3c, regardless of the deviations of *m*^#^ from *m*^*\**^, the adaptation of exoskeleton torque profile leads to total muscles’ effort reduction, which provides strong support for our claim that adaptation of *m* based on feedback from four muscles leads to whole contributor muscles’ effort minimization. In addition, it proves that the suggested sets of mono-articular muscles are sufficient for upper limb exoskeleton torque adaptation. According to Fig.3d-e, four muscles have force reduction while two of them, which are extensor muscles, have force increment. This observation seems evident since flexor muscles are the main contributors to compensate the effect of gravity, and the adaptation rule is trying to minimize the total muscles’ effort cost function, which does not necessarily lead to each individual muscles’ force reduction.

## V. Conclusion and Discussions

In this paper, we presented a novel, simple, and effective human-in-the-loop approach for weight compensation in upper-limb assistive devices. This adaptation rule adapts the exoskeleton torque profile using biofeedback from four mono-articular muscles in order to compensate for the gravitational effects of an unknown object. Our method leads to the whole muscles’ effort reduction, is task-independent, and does not enforce a certain trajectory to the human joints. Hence, the human can have active voluntary behaviors while a portion of the required torque at each joint is compensating by the exoskeleton. The presented method is also analyzed analytically in terms of optimality, stability, and convergence.

Our simulation results support the performance of the proposed method in terms of optimality, stability, and convergence in a generic and realistic model of the human arm in OpenSim, a model with 7-DOF and 50 Hill-type muscles. It is also observed that increasing the frequency/speed of the motions leads to deviations from the unknown mass value; i.e., deviations of equilibrium point from the unknown mass have a linear relation with frequency. However, in all cases, the proposed adaptation rule leads to whole muscles’ effort reduction by feedback from four muscles; a partial observation leads to global optimization. As future work, the goal is to apply this method on a real upper limb exoskeleton device with myoelectric sensors (e.g., EMG) and validate the proposed claim; minimize the whole muscles’ effort using partial observation of four mono-articular muscles.

The proposed adaptation rule for gravity compensation can also be considered a model for mass estimation. This model has the potential to be used as a cognitive model for justifying how humans are interacting with the environment and how the brain and neuromuscular system are combining the sensory muscle information in order to have a quantitative or a qualitative understanding of the environments and object properties. This adaptation rule can also be considered as a deep sensing method for individuals with hand amputations who still have sensory neurons.

## ACKNOWLEDGMENT

The authors would like to thank the University of Tehran for providing support for this work.

## Footnotes

1 In our formulation 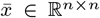 and 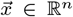 are matrix and vector, respectively while *x*_*j*_ and *x*^*j*^ are both *j*th element of 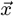.

2 Note that in the rest of the paper, for the sake of simplicity and without loss of generality, we may forbear specifying the argument of the functions, e.g., 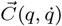 may represent 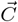.

